# Deamidation of the human eye lens protein γS-crystallin accelerates oxidative aging

**DOI:** 10.1101/2021.06.21.449298

**Authors:** Brenna Norton-Baker, Pedram Mehrabi, Ashley O. Kwok, Kyle W. Roskamp, Marc A. Sprague-Piercy, David von Stetten, R.J. Dwayne Miller, Rachel W. Martin

**Affiliations:** Department of Chemistry, University of California, Irvine 92697-2025, United States; Department for Atomically Resolved Dynamics, Max-Planck-Institute for Structure and Dynamics of Matter, Luruper Chaussee 149, 22761 Hamburg, Germany; Institute for Nanostructure and Solid-State Physics, Universität Hamburg, HARBOR, Luruper Chaussee 149, 22761 Hamburg, Germany; Department of Molecular Biology and Biochemistry, University of California, Irvine 92697-3900, United States; European Molecular Biology Laboratory, Hamburg Unit c/o Deutsches Elektronen-Synchrotron, Hamburg, Germany; Departments of Chemistry and Physics, University of Toronto, 80 St. George Street, Toronto, Ontario, M5S 3H6, Canada

## Abstract

Cataract disease, a clouding of the eye lens due to precipitation of lens proteins, affects millions of people every year worldwide. The proteins that comprise the lens, the crystallins, show extensive post-translational modifications (PTMs) in aged and cataractous lenses, most commonly deamidation and oxidation. Although surface-exposed glutamines and asparagines show the highest rates of deamidation, multiple modifications can accumulate over time in these long-lived proteins, even for buried residues. Both deamidation and oxidation have been shown to promote crystallin aggregation *in vitro*; however, it is not clear precisely how these modified crystallins contribute to insolubilization. Here, we report six novel crystal structures of a major human lens protein, γS-crystallin (γS): one of the wild-type in a monomeric state, and five of deamidated γS variants, ranging from three to nine deamidation sites, after varying degrees of sample aging. Consistent with previous work that focused on single-to triple-site deamidation, the deamidation mutations do not appear to drastically change the fold of γS; however, increasing deamidation leads to accelerated oxidation and disulfide bond formation. Successive addition of deamidated sites progressively destabilized protein structure as evaluated by differential scanning fluorimetry. Light scattering showed the deamidated variants display an increased propensity for aggregation compared to the wild-type protein. The results suggest the deamidated variants are useful as models for accelerated aging; the structural changes observed over time provide support for redox activity of γS-crystallin in the human lens.

**Highlights:**

- Novel structures of cataract-associated variants of human eye lens protein γS-crystallin reported
- Increasing deamidation of γS-crystallin decreases stability and affects aggregation propensity
- Overall fold of γS-crystallin maintained among deamidated and disulfide-bonded variants
- Deamidated γS variants form disulfide bonds more rapidly than wild-type γS
- Potential functional advantage of disulfide bonding in the CXCXC motif proposed

## INTRODUCTION

The eye lens is a unique environment in the human body, hosting some of the most long-lived proteins at extremely high concentrations (Andley, 2007). In order to establish and maintain transparency, the lens cells lose much of their cellular machinery during early development via a combination of autophagy and lipase-mediated destruction of organelles (Morishita and Mizushima, 2016; Morishita et al., 2021). In adulthood, the lens undergoes very little protein turnover (Lynnerup et al., 2008). Crystallins are the predominant protein species in the lens, comprising over 90% of dry mass and reaching concentrations above 400 mg/mL (Bloemendal et al., 2004; Vendra et al., 2016; Wistow, G. J.; Piatigorsky, 1988). Crystallins were originally characterized by their transparency and presence in the “crystalline lens,” and are divided into two superfamilies, α-crystallins and βγ-crystallins (Slingsby et al., 2013). The α-crystallins are related to small heat shock proteins and serve as molecular chaperones that offset aggregation (Horwitz, 1992). The βγ-crystallins are considered structural proteins; they form short-range ordered arrangements and increase the refractive index of the lens, focusing light onto the retina (Serebryany and King, 2014). In order to maintain normal vision, crystallins need to remain stable over the entire human lifetime. However, during aging, modifications accumulate in the eye lens proteins from exposure to reactive oxygen species, ultraviolet light, and lens contamination with non-native molecules and metal ions (Rocha et al., 2021). Due to the presumed loss of structural stability concomitant with these modifications, eye lens proteins aggregate and form large, light-scattering masses in the lens, leading to cataract. Cataract is a leading cause of blindness worldwide, especially in middle- and low-income countries, where surgical intervention is often less accessible (Lee and Afshari, 2017; Liu et al., 2017).

Post-translational modifications (PTMs) of crystallins in the form of deamidation, oxidation, methylation, and truncation have all been observed in aged lenses, with deamidation reported to be one of the most prevalent (Dasari et al., 2009; Lampi et al., 1998; Srivastava et al., 2017; Wilmarth et al., 2006). Deamidation replaces an amide with a carboxylic acid, transforming glutamine or asparagine to glutamic acid or aspartic acid or iso-aspartic acid, respectively, with the possibility of racemization at each site (Geiger and Clarke, 1987; Lampi et al., 2014). Even for surface-exposed residues, deamidation can lead to changes in conformational dynamics and an increase in aggregation propensity (Flaugh et al., 2006; Forsythe et al., 2019; Lampi et al., 2001; Pande et al., 2015; Ray et al., 2016; Takata et al., 2008; Vetter et al., 2020). Additionally, PTMs appear to be interdependent; Vetter *et al*. found that increased deamidation results in an increase in cysteine oxidation and disulfide bonding (Vetter et al., 2020). It is hypothesized that an increase in conformational flexibility in structurally destabilized proteins could lead to solvent exposure of key modifiable residues. Deamidation and disulfide crosslinking are both reported with higher prevalence in cataractous lenses than normal lenses (Hanson et al., 1998; Hooi et al., 2012; Lapko et al., 2002; Takemoto and Boyle, 2000; Wilmarth et al., 2006).

Deamidation has been studied in both β- and γ-crystallins. In β-crystallins, deamidation appeared to alter the dimer structure and led to a less compact fold in βB1 and βA3 (Lampi et al., 2001; Takata et al., 2007). Similar results were found with βB2, with the dimer destabilized by deamidation (Lampi et al., 2006). In the γ-crystallins, γD was destabilized by deamidations at the interface between the two domains of the monomer (Flaugh et al., 2006). However, a number of single-site deamidation variants of surface asparagines on γD did not show any significant changes in structure or stability (Guseman et al., 2021). In contrast, deamidated variants of γS have been reported to have attractive protein-protein interactions, leading to increased propensity for dimerization and aggregation (Forsythe et al., 2019; Pande et al., 2015; Ray et al., 2016; Vetter et al., 2020). Although Vetter *et al*. and Pande *et al*. investigated 3- and 4-site deamidation variants (Pande et al., 2015; Vetter et al., 2020), respectively, most of the previous studies have focused on crystallin variants with only one or two deamidation sites. The present study explores the effects of progressive accumulation of deamidations to assess the tolerance of γS to major surface charge modification, evaluating the type of moderate to extreme deamidation that likely occurs in aged lenses.

Due to the long-lived nature of the crystallins, the lens requires mechanisms of protection and repair to cope with decades of stress. Anti-oxidant compounds, notably glutathione, play vital roles in balancing the redox state of the lens (Michael and Bron, 2011). Crystallin proteins themselves also display adaptations that aid in longevity. Although they have historically been considered purely structural proteins, it has recently been suggested that members of the βγ-crystallin family have chemical functionality in metal binding and oxidation repair in the lens (Roskamp et al., 2020; Serebryany et al., 2018). In particular, γS-crystallin (**Figure 1A**), with its high cysteine content compared to other γ-crystallins, has been proposed to fulfill an oxidoreductase-like role (Roskamp et al., 2020). γS-crystallin contains a cysteine tetrad (C23, C25, C27, C83) in the N-terminal domain (**Figure 1B**), with C23, C25, and C27 exhibiting unusually high solvent exposure. Solvent-exposed cysteines would appear to be disadvantageous for a long-lived protein. Indeed, the dimer of γS-crystallin, formed by the disulfide linkage of C25 and C25 of two γS-crystallin monomers, is less stable and more aggregation-prone than the monomer (Thorn et al., 2019). The functional advantage of these solvent-accessible cysteines in γS-crystallin is still under investigation, although they have been demonstrated to play a critical role in metal interactions. A number of divalent cations, including Zn^2+^, Ni^2+^, Co^2+^ and Cu^2+^, cause aggregation of γS, and the mutation of C23/25/27 dramatically affects this metal-induced aggregation. For Zn^2+^, Ni^2+^, Co^2+^, the mutation of these cysteines to serines prevented aggregation (Roskamp et al., 2019). However, for Cu^2+^, the removal of C23/25/27 increased the susceptibility of γS to Cu^2+^-induced oxidation and aggregation, suggesting that these residues buffer against Cu^2+^-induced oxidative damage (Roskamp et al., 2019, 2020). Additionally, similar solvent-exposed cysteines in γD-crystallin have been shown to undergo dynamic disulfide exchange, transferring disulfide bonds in an oxidoreductase-like role (Serebryany et al., 2018).

**Figure 1.**
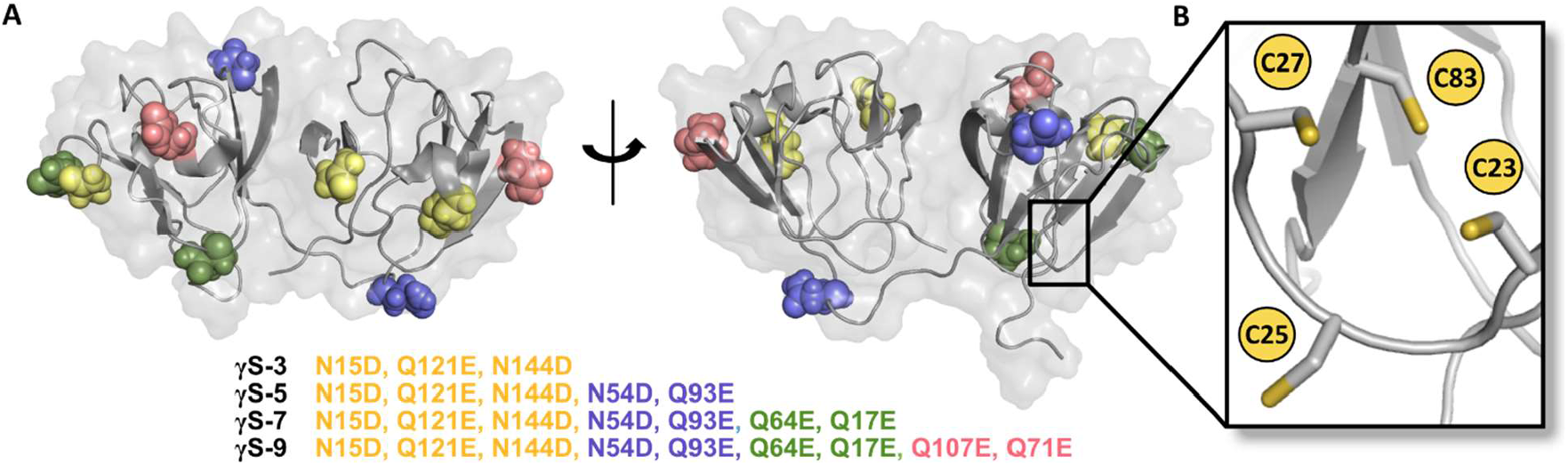
(A) Sites of mutagenesis in the 3-site (γS-3, yellow), 5-site (γS-5, blue), 7-site (γS-7, green), and 9-site deamidation variants (γS-9, pink). (Structure from PDB: 2M3T) (B) Magnified view of the cysteine tetrad located in the N-terminal domain.

In this study, we investigated the relationship between deamidation and disulfide bond formation in γS-crystallin. Crystallins have been shown to collect multiple PTMs; in order to mimic this natural aging process on a compressed timescale, we expressed four deamidated γS-crystallin variants that emulate moderate-to-extreme examples of deamidation, with 3, 5, 7, and 9 sites of deamidation at surface residues. (**Figure 1A**). With each addition of more deamidated sites, the variants demonstrated lower structural stability. Furthermore, the deamidated variants showed an increased aggregation propensity compared to wild-type γS-crystallin (γS-wt). Crystal structures were determined for the full-length, monomeric γS-wt as well as the deamidated variants at different time points in sample age (**Table 1**). Although the deamidation mutations did not cause significant changes to the overall protein fold, these structures revealed that increased deamidation does lead to an increased propensity for disulfide bond formation, indicating accelerated oxidative aging compared to the wild type. An intramolecular disulfide bond between C23 and C27 formed first, with a second dimerizing bond at C25 occurring in increasingly aged samples. Additionally, the structural changes after sample aging give evidence of the role of γS-crystallin as a mediator of oxidation in the eye lens, exchanging deleterious cross-linking disulfide bonds for stable internal disulfide bonds.

**Table 1.** X-Ray Data Collection and Refinement Statistics

|  | **γS-wt** | **γS-3** | **γS-5** | **γS-7** | **γS-9A** | **γS-9B** |
| --- | --- | --- | --- | --- | --- | --- |
| PDB | 7N36 | 7N37 | 7N38 | 7N39 | 7N3A | 7N3B |
| Wavelength (Å) | 0.9762 | 0.9763 | 0.9763 | 0.9763 | 0.9762 | 0.9763 |
| Resolution range | 46.89 - 2.00 | 27.62 - 1.30 | 39.90 - 1.22 | 40.27 - 1.56 | 43.74 - 1.50 | 58.87 - 2.09 |
|  | (2.07 - 2.00) | (1.35 - 1.30) | (1.27 - 1.22) | (1.62 - 1.56) | (1.55 - 1.50) | (2.17 - 2.09) |
| Space group | P2<sub>1</sub> | P2<sub>1</sub>2<sub>1</sub>2<sub>1</sub> | P2<sub>1</sub>2<sub>1</sub>2<sub>1</sub> | P22<sub>1</sub>2<sub>1</sub> | P2<sub>1</sub> | P22<sub>1</sub>2<sub>1</sub> |
| Unit cell |  |  |  |  |  |  |
| a, b, c (Å) | 52.8, 60.6, 56.0 | 29.5, 62.6, 79.6 | 29.3, 62.7, 79.8 | 49.2, 76.6, 94.7 | 29.8, 57.7, 44.9 | 48.7, 75.8, 93.4 |
| $\alpha$, $\beta$, $\gamma$ (°) | 90.0, 117.5, 90.0 | 90.0, 90.0, 90.0 | 90.0, 90.0, 90.0 | 90.0, 90.0, 90.0 | 90.0, 103.3, 90.0 | 90.0, 90.0, 90.0 |
| Total reflections | 146421 (14703) | 473244 (46020) | 528084 (20734) | 682592 (67123) | 160193 (16077) | 270089 (26342) |
| Unique reflections | 20807 (2056) | 36161 (3477) | 41328 (2350) | 51461 (5051) | 23517 (2292) | 20969 (2008) |
| Multiplicity | 7.0 (7.2) | 13.1 (13.2) | 12.8 (8.8) | 13.3 (13.3) | 6.8 (6.9) | 12.9 (13.1) |
| Completeness (%) | 97.42 (97.07) | 97.57 (95.18) | 93.52 (54.14) | 99.61 (99.00) | 98.40 (97.03) | 99.73 (98.14) |
| Mean I/sigma(I) | 16.29 (2.37) | 27.64 (2.00) | 30.24 (2.32) | 27.17 (1.76) | 13.85 (1.24) | 13.98 (1.43) |
| Wilson B-factor | 39.6 | 17.0 | 15.6 | 26.2 | 25.4 | 39.0 |
| R<sub>merge</sub> | 0.06225 (0.7879) | 0.04061 (1.275) | 0.03872 (0.8968) | 0.04674 (1.609) | 0.05397 (1.855) | 0.1236 (1.642) |
| R<sub>meas</sub> | 0.06731 (0.8493) | 0.04234 (1.325) | 0.04035 (0.9517) | 0.04869 (1.674) | 0.05855 (2.004) | 0.1288 (1.708) |
| R<sub>pim</sub> | 0.02531 (0.3145) | 0.01175 (0.3582) | 0.01117 (0.3052) | 0.01345 (0.4546) | 0.02239 (0.752) | 0.03573 (0.4662) |
| CC<sub>1/2</sub> | 0.999 (0.782) | 1.000 (0.795) | 1.000 (0.715) | 0.999 (0.753) | 0.999 (0.734) | 0.998 (0.674) |
| CC* | 1.000 (0.937) | 1.000 (0.941) | 1.000 (0.913) | 1.000 (0.927) | 1.000 (0.920) | 1.000 (0.898) |
| Reflections used in refinement | 20797 (2056) | 36160 (3478) | 41328 (2348) | 51460 (5048) | 23388 (2286) | 20969 (2008) |
| Reflections used for R-free | 962 (101) | 1811 (172) | 2077 (108) | 2650 (256) | 1147 (108) | 1083 (99) |
| R<sub>work</sub> | 0.1891 (0.2478) | 0.1451 (0.2398) | 0.1464 (0.2219) | 0.1934 (0.3036) | 0.1953 (0.4619) | 0.1884 (0.2879) |
| R<sub>free</sub> | 0.2199 (0.3105) | 0.1793 (0.3169) | 0.1792 (0.2817) | 0.2127 (0.3241) | 0.2237 (0.5012) | 0.2396 (0.3641) |
| Number of non-hydrogen atoms | 3030 | 1731 | 1830 | 3442 | 1578 | 3058 |
| macromolecules | 2912 | 1521 | 1588 | 3130 | 1468 | 2935 |
| ligands | - | 29 | 15 | 5 | - | - |
| solvent | 118 | 181 | 227 | 307 | 110 | 123 |
| Protein residues | 348 | 174 | 174 | 348 | 173 | 348 |
| RMS bonds (Å) | 0.006 | 0.014 | 0.010 | 0.006 | 0.008 | 0.007 |
| RMS angles (°) | 0.88 | 1.30 | 1.07 | 0.81 | 0.99 | 0.94 |
| Ramachandran |  |  |  |  |  |  |
| favored (%) | 96.8 | 98.84 | 98.26 | 96.51 | 95.91 | 95.06 |
| allowed (%) | 3.20 | 1.16 | 1.74 | 3.49 | 3.51 | 3.49 |
| outliers (%) | 0 | 0 | 0 | 0 | 0.58 | 1.45 |
| Rotamer outliers (%) | 0.98 | 1.83 | 1.76 | 3.02 | 3.21 | 3.25 |
| Clashscore | 5.67 | 3.62 | 5.43 | 4.26 | 6.00 | 6.90 |
| Average B-factor | 43.59 | 24.73 | 20.92 | 35.75 | 36.02 | 45.70 |
| macromolecules | 43.52 | 23.09 | 19.06 | 34.81 | 35.41 | 45.69 |
| ligands | - | 54.04 | 29.52 | 72.73 | - | - |
| solvent | 45.35 | 33.90 | 33.35 | 44.68 | 44.25 | 46.06 |

## RESULTS

### γS-crystallin deamidated variants are less stable and more aggregation prone than wild-type

We investigated the stability and aggregation propensity of the deamidated variants compared to γS wild-type (γS-wt). Thermal unfolding curves were determined by differential scanning fluorimetry (DSF), and the midpoint of the unfolding temperatures (T_m_) were calculated for each variant (**Figure 2A**). The percent unfolded was calculated via fitting of the averaged fluorescence intensity data (**Supplemental Figure S1**) to a Boltzmann function as described by Wright *et al*. (Wright et al., 2017). The thermal stability progressively decreased with cumulative deamidations from a T_m_ of 76.79 ± 0.06 °C for γS-wt to 73.91 ± 0.07 °C, 72.26 ± 0.01 °C, 71.62 ± 0.02 °C and 70.45 ± 0.04 °C for γS-3, γS-5, γS-7, and γS-9, respectively (**Figure 2B**). A linear relationship emerges from the comparison of the T_m_ and the calculated net charge at neutral pH for each variant, with increased charge leading to decreased stability in these variants. Calculations of net charge were performed with the Prot pi protein tool (https://www.protpi.ch/Calculator/ProteinTool).

**Figure 2.**
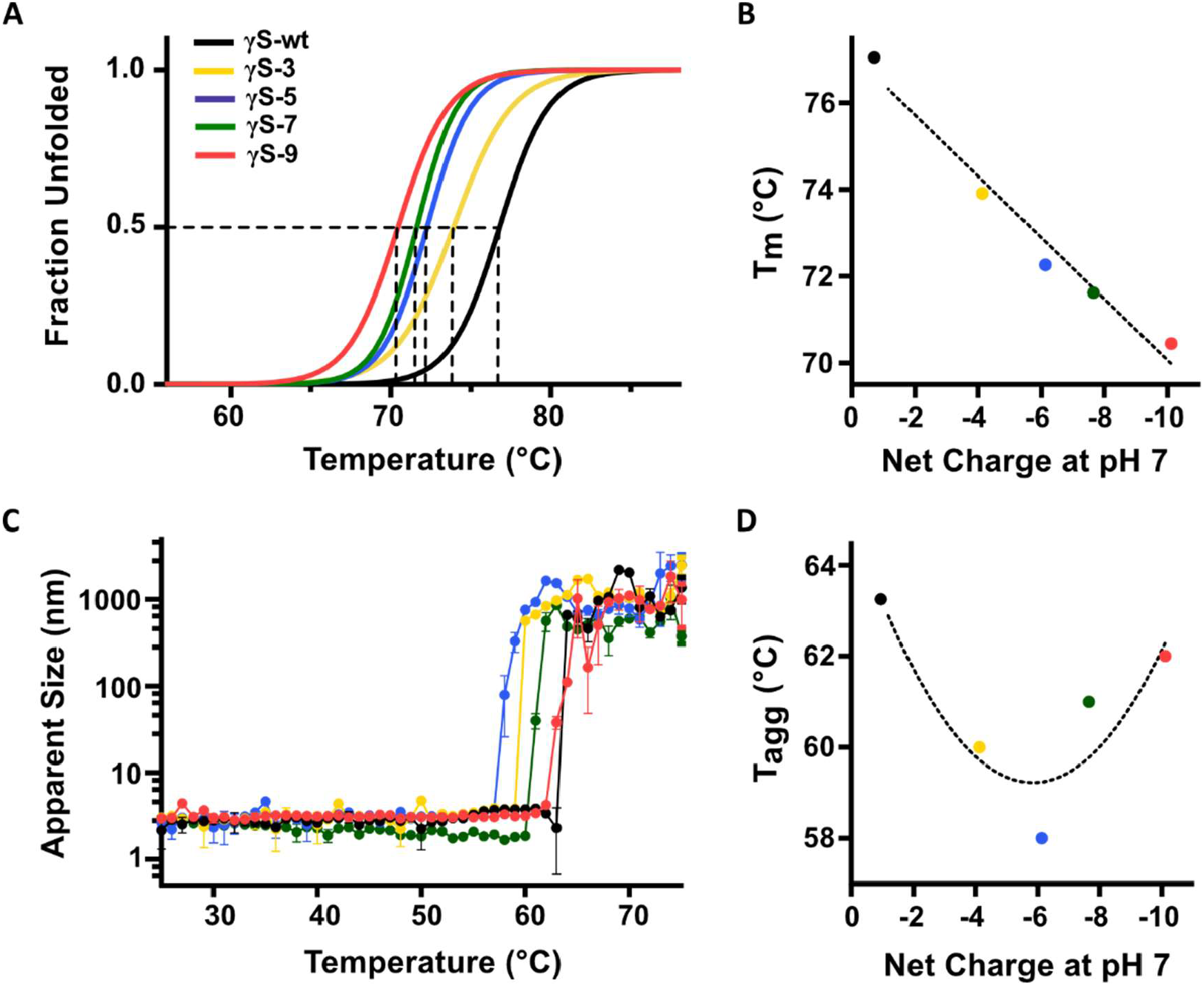
The stability and thermal aggregation of γS-crystallin wild-type (γS-wt, black), 3-site (γS-3, yellow), 5-site (γS-5, blue), 7-site (γS-7, green), and 9-site deamidation variant (γS-9, pink). (A) Differential scanning fluorimetry (DSF) was used to determine the percent unfolded as a function of temperature for γS wild-type and each variant. (B) The midpoint temperature of the thermal unfolding (T_m_) of γS wild-type and each variant plotted against the net charge of the protein at neutral pH. (C) Dynamic light scattering (DLS) measurements of γS wild-type and each variant to monitor thermally induced aggregation. (D) The temperature of aggregation onset (T_agg_) plotted against the net charge of the protein at neutral pH.

Thermally induced aggregation was monitored with dynamic light scattering (DLS). All proteins appeared monomeric at the starting temperature of 25 °C with diameters of 2-4 nm and remained monomeric until the rapid formation of large, insoluble aggregates at the aggregation onset temperature, T_agg_ (**Figure 2C**). The values of T_agg_ were 63 °C for γS-wt and 60 °C, 58 °C, 61 °C, and 62 °C for γS-3, γS-5, γS-7, and γS-9, respectively. All deamidated variants showed a lower T_agg_ compared to γS-wt; however, while T_agg_ for γS-3 and γS-5 decreases with successive deamidations, for γS-7 and γS-9, the trend reverses (**Figure 2D**). We hypothesize that the repulsive effects from the high net charge of γS-7 and γS-9 begin to offset the minor destabilization produced by deamidations, resisting aggregation.

### Crystal structure of monomeric wild-type γS-crystallin

We report the first crystal structure of full-length, monomeric wild-type human γS-crystallin. Descriptive and crystallographic parameters for the structures discussed in this paper are summarized in **Table 1**. The space group was found to be P2_1_ with an asymmetric unit containing two γS-crystallin monomers. In contrast to the previously reported disulfide-bonded dimeric structure, these monomers interact only through non-covalent contacts between the N-terminal domains (Thorn et al., 2019). The structure was solved at a resolution of 2.0 Å and the coordinates have been deposited in the PDB (PDB ID: 7N36). As with other reported γS-crystallin structures, it shows two highly symmetric domains comprising the two double Greek key motifs characteristic of the βγ-crystallins (**Figure 3A**). Structural alignments of the Cα atoms show this novel structure has high similarity with both the γS-wt NMR structure (PDB: 2M3T) and the crystal structure of the C-terminal domain of γS-wt (PDB: 1HA4), with RMSDs of 1.78 and 1.03 Å, respectively, derived by the alignment of chain A of 7N36 with the entire 2M3T structure and chain A of 1HA4 (Kingsley et al., 2013; Purkiss et al., 2002). Likewise, in comparison to disulfide dimerized γS-crystallin structure (PDB: 6FD8), we observe very little structural difference in protein fold, consistent with the analysis by Thorn *et al*., with a Cα RMSD of 0.59 Å, derived from the alignment of chain A of 7N36 and chain A of 6FD8. In both the NMR structure (2M3T) and this novel structure, no disulfide bonds, either intra- or intermolecular, are observed (**Figure 3B**).

**Figure 3.**
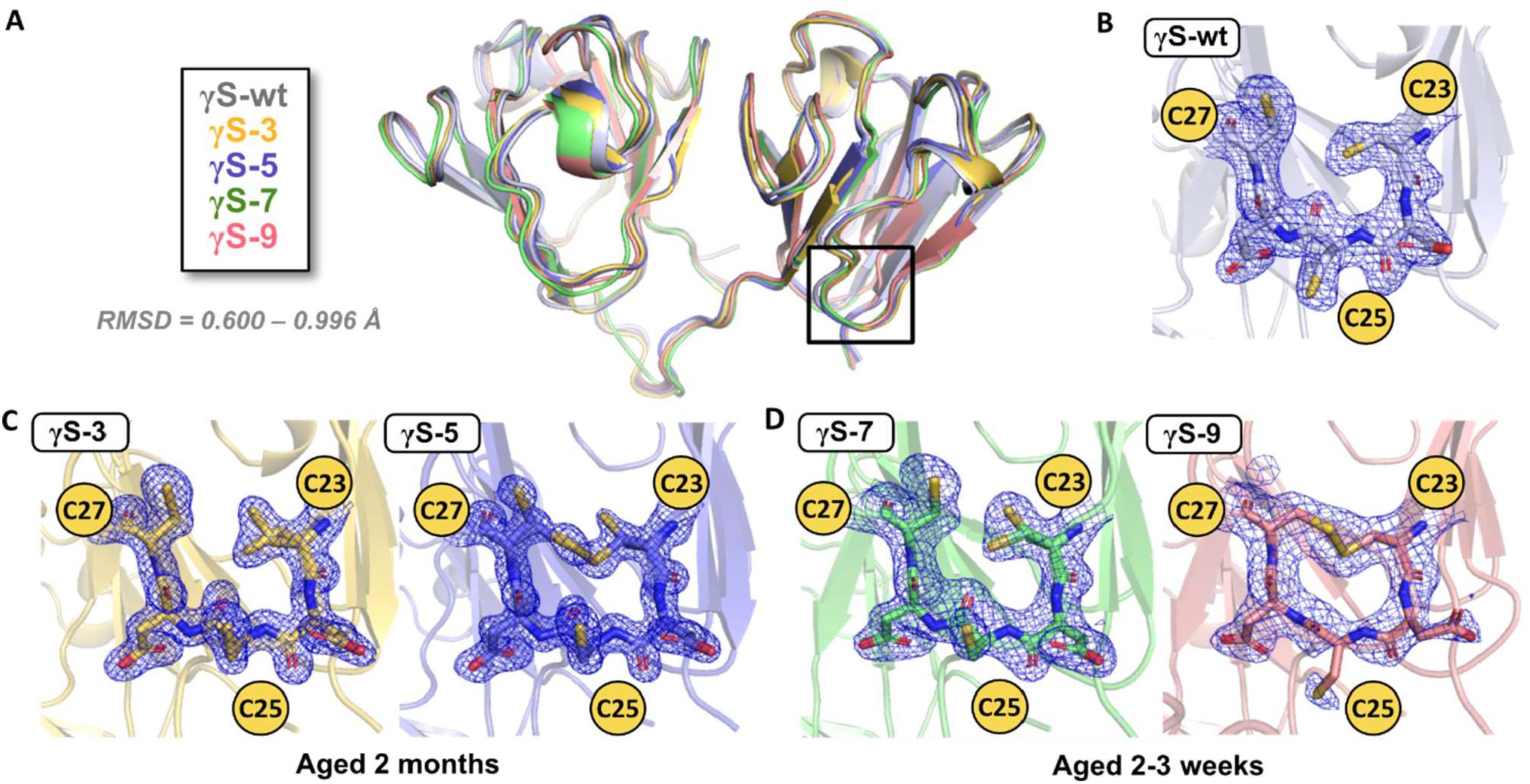
(A) Alignment of the novel structures of γS-crystallin wild-type (γS-wt, PDB: 7N36, gray), 3-site deamidation mutant (γS-3, PDB: 7N37, yellow), 5-site deamidation mutant (γS-5, PDB: 7N38, blue), 7-site deamidation mutant (γS-7, PDB: 7N39, green), and 9-site deamidation mutant (γS-9B, PDB: 7N3B, pink). All structures have Cα RMSDs less than 1 Å compared to γS-wt, PDB: 7N36. Magnified views of the cysteine loop region with 2Fo-Fc (contoured at 1σ) electron density maps for (B) γS-wt aged 1.5 months, (C) γS-3 and γS-5 aged 2 months and (D) γS-7 and γS-9B aged 2-3 weeks.

### Crystal structures of γS-crystallin deamidated variants

Five novel crystal structures have been determined for deamidated variants of γS-crystallin at various stages of sample aging. Sample age is the combined time of storage at 4 °C after resolubilization and time spent in the crystallization plate until the crystals were frozen for diffraction data collection. The γS deamidation variants crystallized in a variety of space groups (**Table 1**). The 3-site deamidation variant (γS-3) and the 5-site deamidation variant (γS-5), both aged for 2 months and grown in the same crystallization solution, were found in space group P2_1_ 2_1_ 2_1_ with a single monomer in the asymmetric unit. The structures for γS-3 and γS-5 were solved at resolution of 1.30 and 1.22 Å, respectively. The refined structures have been deposited in the PDB (PDB ID: 7N37 and 7N38). In both structures, a Mg^2+^ ion was identified, although in different locations. There were no direct interactions between the Mg^2+^ and any residues; interactions occurred through hydrogen bonding with Mg-coordinated waters. γS-crystallin has shown binding and aggregation in the presence of other divalent cations, including Cu^2+^, Zn^2+^, Ni^2+^, and Co^2+^, but shows no such interactions with Mg^2+^ (Roskamp et al., 2019, 2020).

The structures for γS-7 and γS-9B, both aged for 2 – 3 weeks and grown in similar crystallization conditions, were solved at a resolution of 1.56 and 2.09 Å, respectively. They were both found to be in space group P22_1_ 2_1_ with two monomers in the asymmetric unit. The refined structures have been deposited in the PDB (PDB ID: 7N39 and 7N3B). An additional structure was solved of γS-9 from protein aged for 4 – 5 days (γS-9A). γS-9A crystallized in space group P2_1_ with one monomer in the asymmetric unit and was solved at a resolution of 1.50 Å (PDB ID: 7N3A). In general, the deamidated variants grew crystals more readily, in multiple different conditions in the sparse screens, and more reproducibly than γS-wt.

### γS-crystallin deamidation is not significantly detrimental to protein fold

Despite the enhanced crystallization propensity of the variants, the fold of γS-crystallin is remarkably resistant to change after extensive deamidation. Relative to the new γS-wt crystal structure (PDB: 7N36), the backbone fold is maintained in all deamidation variants, even for the highly deamidated γS-9. The Cα RMSD relative to the wild-type structure does not exceed 1 Å for any deamidated variant (**Figure 3A**). Additionally, the intramolecular disulfide bond observed in the γS-9B crystal structure does not seem to significantly affect protein fold, with a Cα RMSD of 1.19 Å in comparison to chain A of γS-9B to the γS-9A structure.

### Deamidated γS-crystallin variants have a higher propensity for disulfide formation

Comparisons of different crystal structures can be challenging, as many factors related to crystallization or diffraction data collection, rather than the protein structure, can have substantial effects. To this end, the crystals directly compared here (γS-3 vs γS-5 and γS-7 vs. γS-9B) are from protein samples that were grown in identical or highly similar crystallization conditions and formed crystals in the same space group, with similar unit cells, and of comparable sizes. The data collection strategies were kept uniform so that radiation dose would also be consistent between samples.

Comparing crystal structures from samples aged for similar periods, we observe an increase in disulfide bond formation in variants with more deamidation. After aging for 2 months, the γS-3 structure has fully reduced cysteines (**Figure 3C**). In contrast, the γS-5 structure shows a mixed state for C23 and C27, with both reduced and disulfide-bonded species present (**Figure 3C**). Occupancy refinement in Phenix suggests that the disulfide-bonded configuration accounts for ∼50-70%, with a range estimated due to the different occupancy calculation for the two cysteines, C23 and C27. The same trend is observed in samples aged 2-3 weeks, γS-7 and γS-9B. The γS-7 structure shows the cysteines to be fully reduced, while the γS-9B structure shows C23 and C27 are fully oxidized, forming an intramolecular disulfide bond (**Figure 3D**). The fourth cysteine in the tetrad, C83, is not shown in these maps, but in all cases it appears to be fully reduced. This may indicate that C83 is not amenable to disulfide bonding or that the C83 disulfide-bonded structure is unfavorable for crystallization, possibly due to large structural disruption from this bond.

### γS accumulates disulfide bonds during sample aging

Because no significant structural disruption to protein fold is observed in the deamidated variants, we propose that they may serve as more crystallizable models for the wild-type protein to gain atomic and near-atomic resolution structures at varying time points. Structural changes over time in the deamidated variants may give insight into the aging behavior of the wild-type protein, but in an accelerated timeframe. In this study, γS-9 was crystallized at two time points of sample age: several days and several weeks. Recently, we reported another crystal structure of γS-9 that was aged for several months prior to crystallization in a silicon chip. That structure was solved via serial crystallography on >1000 microcrystals to a resolution of 3.0 Å (referred to here as γS-9C, PDB: 7NJE) (Norton-Baker *et al*., 2021). The comparison of these three structures allows us to observe the change in cysteine oxidation over time (**Figure 4**). In the earliest time point structure, γS-9A, the protein is monomeric and all three cysteines appear to be mostly reduced, with minor partial occupancy of a C23-C27 disulfide bond (**Figure 4A**). After several weeks, as discussed above, γS-9B has a full intramolecular disulfide between C23 and C27 and the protein remains monomeric (**Figure 4B**). After several months, γS-9C contains a second disulfide bond between C25 and C25 of another γS-9, forming a dimer (**Figure 4C**). Interestingly, the orientation of C25 shifts after the formation of the C23-C27 disulfide bond. In γS-9A, the side chain of C25 projects out from the ring and has a high calculated solvent exposed surface area (SASA) of 85 Å^2^. Similarly, γS-wt shows fully reduced cysteines with a SASA of 82 Å^2^ calculated for C25. In γS-9B, after the formation of the C23-C27 disulfide, C25 flips and the SASA is lowered to 30 Å^2^. SASA values were calculated by AREAIMOL in the CCP4 package (Lee and Richards, 1971; Winn et al., 2011).

**Figure 4.**
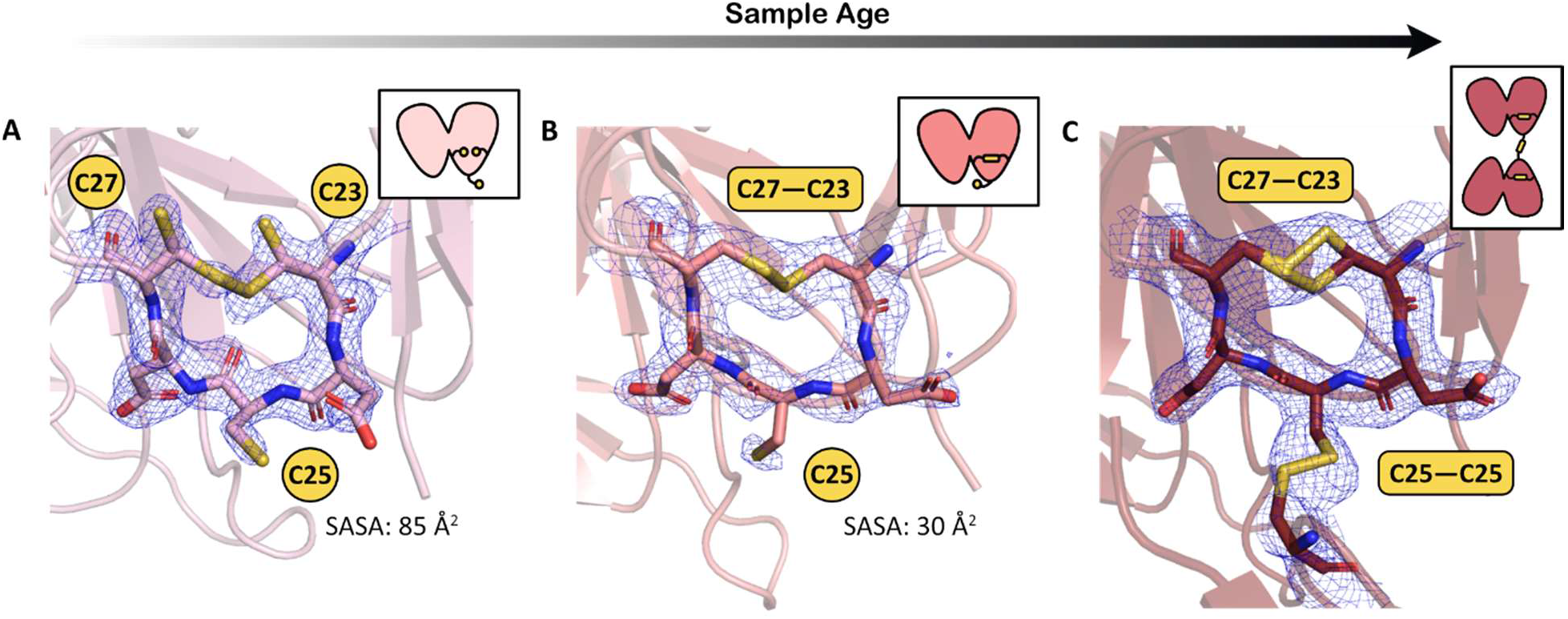
Magnified views of the cysteine loop region with 2Fo-Fc (contoured at 1σ) electron density maps for (A) γS-9 aged for several days (γS-9A, PDB: 7N3A), (B) γS-9 aged for several weeks (γS-9B, PDB: 7N3B), and (C) γS-9 aged for several months (γS-9C, PDB: 7NJE). Insets show a cartoon depiction of the γS-9 structures indicating their oligomeric states (either monomeric or dimeric) with intramolecular and intermolecular disulfide bonds in the cysteine loop shown. The solvent accessible surface area (SASA) for C25 is lower after disulfide bond formation between C23 and C27.

## DISCUSSION

### Deamidation accelerates oxidative aging

In comparing structures of samples of γS-wt and deamidation variants aged for time periods ranging from several days to several months, we observe an increased propensity for disulfide bonding with increased deamidation. The duration of sample aging of γS-wt, γS-3 and γS-5 was similar, between 1.5 – 2 months; however, only γS-5 shows partial occupancy of disulfide bonded C23-C27. At an earlier aging timepoint, γS-7 and γS-9 can be compared after several weeks aging; γS-7 remains fully reduced, while γS-9 shows a full disulfide bond between C23-C27. Based on this trend, we propose that deamidation accelerates the process of oxidative aging, possibly by increasing conformational flexibility, which then increases solvent exposure of the cysteine residues. Other studies have linked deamidation with an increase in conformational dynamics and disulfide bonding. Forsythe *et al*. showed that deamidation of two surface Asn residues in the N-terminal domain of γS caused long-range disruption of conformational dynamics in the C-terminal domain (Forsythe et al., 2019). Their work suggested that deamidation induced both local and global transient unfolding. Vetter *el al*. followed up on this study, demonstrating that with three deamidation mutations, γS became progressively more susceptible to oxidation with each mutation (Vetter et al., 2020).

The mutated residues (glutamine to glutamic acid and asparagine to aspartic acid) are structurally highly similar; therefore, the difference in charge after mutation is the likely basis for the altered structural dynamics. Upon deamidation, a negative charge is introduced at each deamidated site as the carboxylic acid moiety is ionized at the relevant pH values. These charges not only lower the pI of the protein significantly, but also induce regions of high negative charge on the surface of the protein (**Supplemental Figure S2**). At neutral pH, γS-wt has an expected net charge of -1. With each additional deamidation mutation, the net charge decreases, reaching -10 for γS-9. A complex picture emerges from the evaluation of the benefits and drawbacks of introducing negative charges. Chong *et al*. proposed that additional negative charges increase the solubility for proteins that already have a net negative charge (Chong and Ham, 2014). However, the introduction of charged sites can also lead to an increase in fluctuating conformational transitions, or transient misfolding, which can drive aggregation through exposure of hydrophobic portions of the protein (Chong and Ham, 2015).

Our findings support the hypothesis of the competing role between net charge and conformational dynamics in governing aggregation propensity. As the deamidated variants gained negative charge, their structural stability decreased, as indicated by the observed unfolding temperatures (**Figure 2B**). Lower stability likely promotes transient misfolding, prompting an earlier onset of thermally induced aggregation in moderately charged variants (γS-3 and γS-5); however, as the proteins reached high negative charge (γS-7 and γS-9), the greater electrostatic repulsion between molecules raises the energic barrier to aggregation (**Figure 2D**). Further studies into the local and global dynamic effects of charged residues are warranted.

### γS serves as an oxidation sink in the lens

Previously, disulfide bonding in γS was thought to be solely a detrimental byproduct of oxidation. Recent work and the results of this study suggest that it may also be a functional adaptation that helps to sustain the redox balance in the lens (Roskamp et al., 2020; Serebryany et al., 2018). Here, we demonstrate that deamidation accelerates the oxidation of the C23, C25, and C27. Due to the unchanged global fold across the variants characterized in this study, we propose that we may extrapolate the behavior of the deamidated variants to the wild-type protein observed over a longer time period.

We suggest that the positioning of a highly solvent-exposed cysteine, C25, may be advantageous in γS due to its proximity to other cysteines that can form a stable internal disulfide bond. A possible mechanism for γS oxidoreductase activity emerges from the observations of the crystal structures determined in this study (**Figure 5**). Fully reduced, monomeric γS (**1**) contains the highly solvent-exposed C25, predisposing it to dimerization with a second molecule of γS. Although this schematic diagrams the *in vitro* process with only γS available for crosslinking, in the lens this may also occur with a different crystallin species. A transient dimer (**2**) is formed with a crosslinking disulfide bond between C25 and a C25 of another crystallin monomer. This dimerizing bond exchanges with other nearby cysteines, forming an intramolecular disulfide bond between C23-C27 (**3**). This process removes a crosslinking disulfide bond and releases a recovered, reduced monomer of crystallin. After the formation of the C23-C27 bond, C25 occupies an orientation with reduced solvent exposure, as evidenced by the crystal structure of γS-9B. Further oxidation eventually leads to the formation of a fully oxidized and aggregation-prone dimer, but this conversion may be slowed by the reduced solvent exposure of C25. The C23-C27 bond is apparently not highly disruptive to the protein fold in this region, indicating that this bond might serve to rescue γS or other crystallins from other, more deleterious disulfide bonds. If this hypothesis is correct, γS plays not only a structural role, but also a chemical role in the lens, acting as a final buffer against oxidative damage, even after depletion of glutathione.

**Figure 5.**
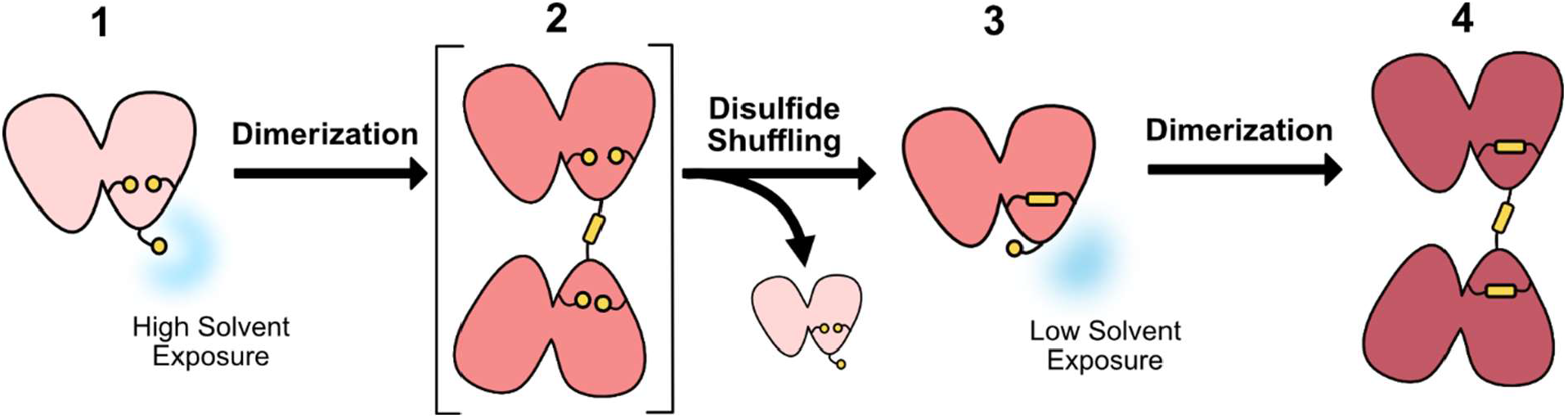
Potential mechanism for the role of γS-crystallin as an oxidation sink in the eye lens. Monomeric, reduced γS-crystallin (**1**) with a highly solvent exposed C25 accepts an intermolecular disulfide bond from another γS-crystallin or another oxidized crystallin in the eye lens. This partially oxidized dimer (**2**) undergoes disulfide shuffling and the disulfide bond exchanges to C23-C27 forming an intramolecular disulfide bonded monomer (**3**) and releasing a recovered, reduced crystallin monomer. The reduced solvent exposure of C25 in the single disulfide bond intermediate (**3**) slows the formation of the fully oxidized disulfide-bonded dimer (**4**) that has been shown to be prone to aggregation.

In other proteins with disulfide reactivity, a CXXC motif has been reported, notably in thioredoxins and disulfide isomerases. The XX is often proline and glycine, both of which support a tight turn and the formation of a disulfide between the two cysteines (Oka et al., 2015). The same activity is recapitulated with the CXC motif, where X is usually glycine (Wilkinson and Gilbert, 2004). The activity of a CXC motif has been previously reported for human γD-crystallin, in a study showing disulfide exchange between wild-type γD and an aggregation prone variant, W42Q (Serebryany et al., 2018). In this case, X is serine, a highly flexible amino acid that is often present in flexible linker regions (Huang and Nau, 2003; Van Rosmalen et al., 2017); however, the disulfide bond was still postulated to cause conformational strain in the C-terminal domain. Upon disulfide exchange between the wild-type and W42Q variant, γS-W42Q was more destabilized by the disulfide and more rapidly aggregated out of solution (Serebryany et al., 2018). γS contains a similar, extended CXCXC motif. However, instead of glycine or proline that allow for tight turns, here the cysteines are separated by aspartic acid residues. As the structures reported here show, the favored disulfide forms between the first and last C in the motif (C23 and C27); the 3 separating residues likely allow for a less strained disulfide bond formation. Aspartic acid is, however, among the most flexible amino acids (Huang and Nau, 2003), lessening any structural disruption as well as supporting disulfide exchange between C25 and C23/27.

The presence of aspartic acid in this motif raises additional questions; is there a fitness benefit from proximal acidic residues? Repulsive electrostatic interactions between deprotonated cysteine and neighboring negatively charged residues have been suggested to destabilize the thiolate state and thereby raise cysteine p*K*_a_ (Awoonor-Williams and Rowley, 2016). As thiolate is a much stronger nucleophile than the protonated thiol, an increase in cysteine p*K*_a_ would likely be inhibitory to thiol-disulfide exchange (Bulaj et al., 1998; Nagy, 2013). Free cysteine in solution has a reported thiol p*K*_a_ of 8.6 and most noncatalytic cysteine p*K*_a_’s fall within 7.4 – 9.1 (Awoonor-Williams and Rowley, 2016); however, dramatic shifts can been seen for catalytic cysteines, with p*K*_a_ values as low as 2.88 reported (Pinitglang et al., 1997). The predicted p*K*_a_ for C25 in γS is 9.35, calculated with PROPKA using the γS-wt crystal structure, 7N36 (Olsson et al., 2011; Søndergaard et al., 2011). C25 is therefore predicted to be slightly more basic than glutathione, the major antioxidant in the lens with a reported thiol p*K*_a_ of 8.92 (Schafer and Buettner, 2001). In addition to similar p*K*_a_ values, C25 in the CDCDC motif of γS and the structure of glutathione also share structural similarities. Glutathione is a tripeptide (L-γ-glutamyl-L-cysteinyl-glycine) with the amide bond formed with the γ-carboxyl rather than the α-carboxyl of glutamic acid. Like C25 in γS, the cysteine in glutathione is flanked by adjacent carboxylates. We plan to further investigate the role of γS and its interplay with glutathione in maintaining redox homeostasis in the eye lens.

### Deamidation as a basis for rational mutagenesis for crystallization

On a more methodological note, deamidation may serve as a new target for rational mutagenesis to promote crystallization. Excellent work has been done in this area, with much of the effort focused on surface entropy reduction (SER) (Goldschmidt et al., 2007). In SER, surface residues with high conformational energy, such as Lys and Glu, are mutated to Ala or other small amino acids. This method is often successful but may sometimes lead to undesired structural changes. The deamidated variants reported here did not appear to change the overall fold of the protein but did more readily and reproducibly form crystals. For challenging crystallization targets, deamidation of surface Gln and Asn to Glu and Asp may provide less disruption to protein fold than other mutations, as the side chains are highly similar in size.

In conclusion, crystallins are among a unique class of extremely long-lived proteins (ELLPs). The half-lives of ELLPs far exceed those of average proteins, persisting for years or even the entire lifetime of an organism (Toyama et al., 2013). In metabolically inactive cells such as the eye lens, the cells’ resources to safeguard against protein degradation and aggregation are finite. Multiple and redundant mechanisms for maintaining lens transparency are likely necessary and the γ-crystallins appear to have a multidimensional role in the lens. γS-crystallin demonstrates resistance to significant structural change from deamidation mutations; however, these mutations predispose the protein to accumulate oxidative modifications. Furthermore, although glutathione is the main species responsible for maintaining redox balance in the lens, here we provide further evidence that γS-crystallin may also aid in reducing oxidized species by shuffling disulfide bonds internally to non-crosslinking positions. By this mechanism, γS may act as a last line of defense against oxidation. The proteins of the eye lens appear to employ methods to both counteract and exploit PTMs; the full role of crystallins and other lens molecules in slowing the irreversible aggregation that leads to cataract remains to be fully explored.

## METHODS

### γS-crystallin expression and purification

Sites for deamidation were selected based on mass spectrometry data reported by Lapko *et al*. (Lapko et al., 2002), prioritizing sites with the highest reported incidence of deamidation and those with the highest calculated solvent-accessible surface area. Progressively more sites were mutated, starting with N14D, Q121E, N144D (3-site variant, γS-3), then N54D and Q93E (5-site variant, γS-5), then Q64E and Q17E (7-site variant, γS-7), and finally Q107E and Q71E (9-site variant, γS-9) (**Figure 1A**). Site-specific mutagenesis PCR was used to create the deamidated variants from the wild-type construct with an N-terminal 6x-His tag and a tobacco etch virus (TEV) protease cleavage sequence. The oligonucleotides used in the mutagenesis are listed in **Table S1**. Expression and purification of wild-type γS-crystallin (γS-wt) and the deamidated variants was performed as described previously (Brubaker et al., 2011). The constructs in pET28a(+) vector were used to transform *Escherichia coli* Rosetta (DE3) cells. Overexpression was achieved using Studier’s autoinduction method for 1 h at 37 °C followed by 20−24 h at 25 °C incubation (Studier, 2005). The cells were lysed by sonication and the supernatant loaded onto a Ni-NTA column (Applied Biosystems, Foster City, CA, USA). An imidazole gradient was used to elute the tagged protein, then TEV protease (produced in-house) was added to cleave the 6x-His tag. Reapplication to the Ni-NTA column separated the TEV protease. The samples were concentrated and loaded on to a HiLoad 16/ 600 Superdex 75 pg column (GE Healthcare Life Sciences, Piscataway, NJ, USA). For biophysical measurements, the proteins were used within days after initial purification. For crystallization, the purified proteins were lyophilized for storage at -80 °C.

### Differential scanning fluorimetry

Differential scanning fluorimetry (DSF) analysis was performed in a Stratagene Mx3005P RT-PCR instrument (Agilent, Santa Clara, CA, USA) in thin-walled 96-well PCR plates with transparent Microseal® ‘B’ seals (Bio-Rad, Hercules, California, USA). The temperature ramp was between 25 °C and 94 °C with a 0.5 °C increase per cycle for 140 cycles and a cycle duration of 30 seconds. Midpoint of the unfolding temperatures (T_m_) were calculated as described by Wright *et al*. (Wright et al., 2017). Each well contained 0.5 mg/ml protein and 2.5X SYPRO orange dye (Sigma, St. Louis, MO, USA) in 10 mM HEPES pH 7.0, 50 mM NaCl, 0.05% sodium azide.

### Dynamic light scattering

Thermally-induced aggregation was assessed with dynamic light scattering (DLS) using a Zetasizer Nano-ZS (Malvern Analytical, Malvern, United Kingdom). Protein concentration was 1 mg/ml in 10 mM HEPES pH 7.0, 50 mM NaCl, 0.05% sodium azide. At each temperature, the sample was equilibrated for 120 seconds before the scattering measurements were performed in triplicate. Data was processed using the Zetasizer software. The autocorrelation function was used to derive the intensity of scattering as a function of particle size. A number distribution is then calculated to estimate the relative concentrations of particles of different sizes, accounting for their proportional light scattering. The reported apparent size is the number mean.

### γS-crystallin crystallization, data collection, and structure refinement

Crystallization conditions were found via sparse matrix screening. Crystallization occurred at 20 °C and the protein:precipitant ratio was 1:1. For all protein samples, the lyophilized powder was resolubilized and buffer exchanged into 50 mM HEPES pH 6.8 with 5 mM DTT. The addition of DTT allowed for a uniform starting condition for all samples with cysteines reduced.

For γS-wt, γS-3, and γS-5, the samples were aged overnight at 4 °C before crystallization. For γS-wt, crystals were grown in the Morpheus screen (Molecular Dimensions, Maumee, OH, USA) condition G5 (0.1 M Carboxylic Acids Mix, 0.1 M Buffer System 2 pH 7.5, 30 % v/v Precipitant Mix 1). The Carboxylic Acids Mix contains 0.2 M sodium formate, 0.2 M ammonium acetate, 0.2 M trisodium citrate, 0.2 M sodium potassium l-tartrate, and 0.2 M sodium oxamate. Buffer system 3 contains 0.1 M MOPS/HEPES-Na pH 7.5. Precipitant Mix 1 contains 20% w/v PEG 20 000 and 40% v/v PEG MME 550 (Gorrec, 2009). Protein concentration was 12 mg/mL. Crystals appeared between 14-28 days and were harvested after 1.5 months. For γS-3 and γS-5, crystals were grown in the Hampton Index screen (Hampton Research, Aliso Viejo, CA, USA) condition G10 (0.2 M MgCl_2_, 0.1 M BIS-TRIS pH 5.5, 25 % w/v Polyethylene glycol 3350). Protein concentration was 18 mg/mL. Crystals appeared between 14-28 days and were harvested after 2 months.

For γS-7, the protein sample was aged at 4 °C for 1-2 weeks before being crystallized in 0.1 M sodium acetate pH 5.0, 0.1 M ammonium sulfate, 28% PEG 3350. Crystals appeared after 1 day and were harvested after 1 week. For γS-9A, the protein sample was aged overnight at 4 °C before being crystallized in 0.1 M sodium acetate pH 5.45, 21% PEG 3350. Crystals appeared after 1 day and were harvested after 2-3 days. For γS-9B, the protein sample was aged at 4 °C for 1-2 weeks before being crystallized in 0.1 M sodium acetate pH 5.45, 0.1 M ammonium sulfate, 21% PEG 3350. Crystals appeared after 1 day and were harvested after 1 week.

Crystals were cryoprotected with Paratone-N (Hampton Research, Aliso Viejo, CA, USA) and flash-frozen by plunging into liquid nitrogen. X-ray diffraction data were collected at both the P13 or P14 beamline operated by the European Molecular Biology Laboratory (EMBL) at the PETRA-III synchrotron at DESY, Hamburg, Germany (https://www.embl-hamburg.de/services/mx/P13/ or https://www.embl-hamburg.de/services/mx/P14/). Diffraction was recorded on a PILATUS 6M or EIGER 16M detector. Diffraction data was processed using XDS (Kabsch, 2010). Phasing was performed using Phaser molecular replacement with chain A of PDB model 6FD8 as a search model (McCoy et al., 2007). Refinement was performed using the Phenix software suite with iterative manual improvement in Coot (Emsley et al., 2010; Liebschner et al., 2019). Structure images were generated in PyMOL (Schrödinger LLC, 2015).

## ACKNOWLEDGMENTS

This investigation was supported by NIH grant 2R01EY021514 to R.W.M. and CIFAR (R.W.M and R.J.D.M). B.N.-B. acknowledges support from the NSF GRFP and the Fulbright Fellowship. P.M. acknowledges support from the Joachim Herz Stiftung Add-on Fellowship and the Deutsche Forschungsgemeinschaft via grant No. 451079909. M.A.S.-P. acknowledges support from the HHMI Gilliam Fellowship. K.W.R. acknowledges support from the MAPS Training Grant, NSF DGE-1633631. The authors thank Dmitry Fishman and Felix Grun and Ben Katz for excellent management of the UCI Laser Spectroscopy Laboratories and Mass Spectrometry Facility, respectively.

## AUTHOR CONTRIBUTIONS

B.N.-B, K.W.R, and R.W.M designed the protein constructs and planned the biophysical experiments. B.N-B., P.M, R.J.D.M., and R.W.M. designed the crystallographic studies. B.N.-B, A.O.K, and M.A.S.-P prepared the protein. B.N.-B, P.M., and D.v.S. collected and processed crystallography data. B.N.-B performed the DSF and DLS experiments. B.N.-B and R.W.M. wrote the manuscript. All authors edited the manuscript.

