## Supplementary Material for "Deamidation of the human eye lens protein γS-crystallin accelerates oxidative aging"

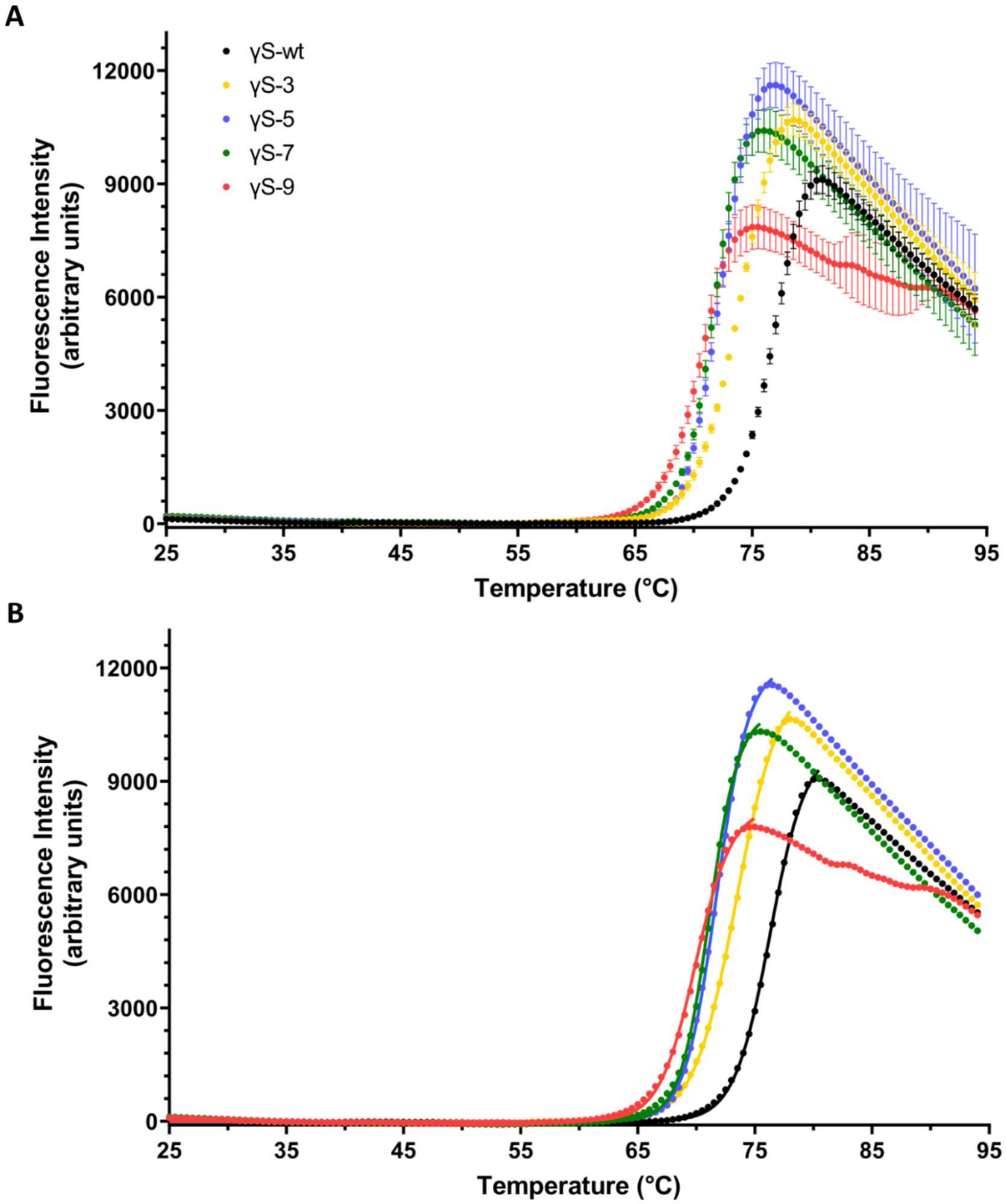


**Figure S1.** (A) The average fluorescence intensity from three replicates measured by differential scanning fluorimetry (DSF) for γS-crystallin wild-type (γS-wt, black), 3-site (γS-3, yellow), 5-site (γS-5, blue), 7-site (γS-7, green), and 9-site deamidation variant (γS-9, pink). (B) The average fluorescence intensity for each protein with overlaid solid lines showing the Boltzmann fit curves calculated for each data set. Only the region of the DSF data up to the fluorescence maxima are used for the Boltzmann fit as the fluorescence diminishes after this point due to protein aggregation.

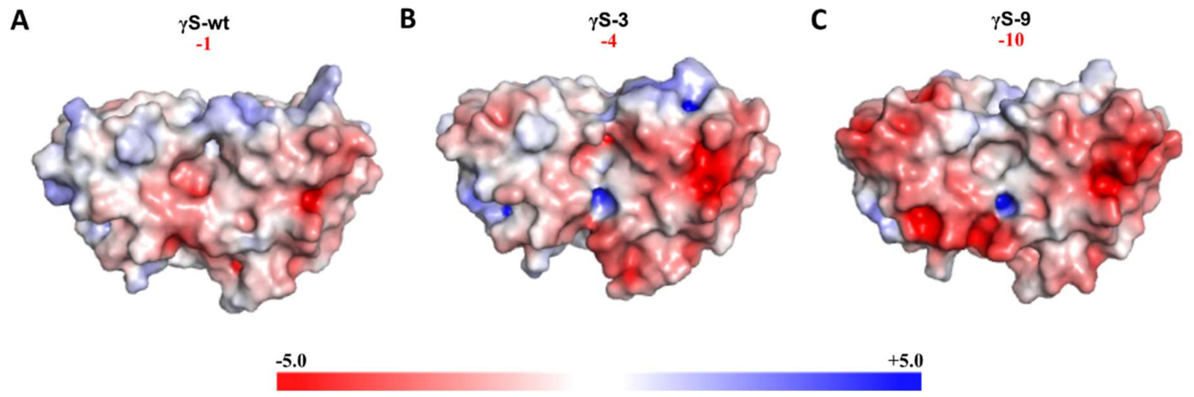


**Figure S2.** Adaptive Poisson-Boltzmann Solver (APBS) electrostatics calculation was used to generate surface charge maps in PyMol for (A) γS-wt PDB:7N36, (B) γS-3 PDB:7N37, and (C) γS-9B PDB:7N3B showing regions of high negative charge with increasing deamidation.

**Codon-optimized γS-wt sequence with NcoI and XhoI restriction sites:**

CCATGGGACATCACCATCACCATCACGAAAATCTTTATTTTCAAGGCAGCAAAACGGGCACCAAAATCACGTTC
TACGAAGACAAAAATTTCCAAGGTCGCCGTTACGACTGTGACTGCGACTGTGCGGACTTCCACACATACCTGTC
CCGTTGCAACTCTATCAAAGTTGAAGGCGGCACCTGGGCTGTATACGAACGCCCGAACTTCGCTGGCTACATGT
ACATCCTTCCGCAGGGCGAATACCCGGAATACCAGCGCTGGATGGGTCTGAACGACCGTCTGTCGTCTTGCCGC
GCTGTACATCTGCCGTCCGGCGGTCAGTACAAAATCCAGATCTTCGAAAAAGGCGACTTCTCCGGCCAGATGTA
CGAAACCACCGAAGACTGCCCGTCTATCATGGAACAATTCCACATGCGCGAAATCCACTCTTGTAAAGTTCTGG
AAGGCGTATGGATCTTCTACGAACTGCCGAACTACCGCGGTCGCCAGTACCTGCTGGACAAAAAAGAATACCGC
AAACCGATCGACTGGGGCGCGGCCTCTCCGGCTGTACAGAGCTTCCGTCGTATCGTTGAATAATAACTCGAG

**Table S1.** Oligonucleotides for PCR mutagenesis. A mistranslation of Y11 was identified in the deamidated variants and was corrected by the application of primers *codonY11_fwd* to γS-3 and γS-5 and *codonY11_fwd_2* to γS-7 and γS-9 with *codonY11_rev* used for all deamidated variants.

| Primer | Sequence (5' to 3') |
|---|---|
| N15_fwd | GAC AAA GAT TTC CAA GGT CGC |
| N15_rev | GCG ACC TTG GAA ATC TTT GTC |
| Q121_fwd | ATC ATG GAA GAA TTC CAC ATG |
| Q121_rev | CAT GTG GAA TTC TTC CAT GAT |
| N144_fwd | TAC GAA CTG CCG GAC TAC |
| N144_rev | GTA GTC CGG CAG TTC GTA |
| N54D_fwd | CCG GAC TTC GCT GGC TAC ATG TAC ATC CTT CCG |
| N54D_rev | GCG TTC GTA TAC AGC CCA GGT GCC GCC TTC |
| Q93E_fwd | CGT CCG GCG GTG AGT ACA AAA TCC AGA TCT TCG AAA AAG GCG |
| Q93E_rev | GCA GAT GTA CAG CGC GGC AAG ACG ACA GAC GG |
| Q64E_fwd | CTT CCG GAG GGC GAA TAC CCG GAA TAC CAG C |
| Q64E_rev | GAT GTA CAT GTA GCC AGC GAA GTC CGG GCG TTC GTA TAC |
| Q17E_fwd | GAA GAC AAA GAT TTC GAA GGT CGC CGT TAC GAC TGT GAC TG |
| Q17E_rev | GTA GAA CGT GAT TTT GGT GCC CGT TTT GCT GCC TTG |
| Q107E_fwd | GGC GAG ATG TAC GAA ACC ACC GAA GAC TGC CCG TCT ATC |
| Q107E_rev | GGA GAA GTC GCC TTT TTC GAA GAT CTG GAT TTT GTA CTC ACC GCC G |
| Q71E_fwd | GGA ATA CGA GCG CTG GAT GGG TCT GAA CGA CCG TCT GTC G |
| Q71E_rev | GGG TAT TCG CCC TCC GGA AGG ATG TAC ATG TAG CCA GCG |
| codonY11_fwd | CAA AGA CTT CCA AGG TCG CCG TTA C |
| codonY11_fwd_2 | CAA AGA CTT CGA AGG TCG CCG TTA C |
| codonY11_rev | TCT TCA TAG AAC GTG ATT TTG GTG CC |
